# DIO-SPOTlight Transgenic Mouse to Functionally Monitor Protein Synthesis Regulated by the Integrated Stress Response

**DOI:** 10.1101/2024.10.14.618312

**Authors:** Matthew L. Oliver, Zachary F. Caffall, Callie B. Eatman, Timothy D. Faw, Nicole Calakos

## Abstract

The integrated stress response (ISR) is a core pathway for maintaining cellular proteostasis and a key regulator of translation in processes beyond the cellular response to stress. For example, the ISR regulates developmental axonogenesis, learning and memory, and synaptic plasticity in the brain. One barrier to uncovering ISR roles in health and disease is the challenge of monitoring its activity. The transient nature of regulatory phosphorylation events and lack of transgenic ISR reporter mouse lines make visually capturing the molecular hallmarks of ISR activation in specific cell types especially difficult. We recently developed the SPOTlight (<u>S</u>elective <u>P</u>hospho-eIF2α <u>O</u>pen reading frame <u>T</u>racking light) reporter, which uniquely provides a readout of the functional state of protein synthesis initiation dynamics that are regulated by the ISR. Here, we report the generation of a transgenic mouse line with Cre-dependent expression of SPOTlight. This resource enables selective visualization of ISR-regulated functional activity across genetically defined cell populations body-wide. Using a pan-neuronal Cre line (*Nestin*-Cre), we demonstrate the reporter’s performance and applications for cell-specific discovery, live tissue assessments and quantitative comparisons across broad physical space. We also specifically investigated the extent to which the property of steady-state basal ISR activation, recently described in dorsal striatal cholinergic interneurons, extends to other classes of cholinergic neurons and provide a CNS-wide atlas of SPOTlight activity in these cells. The DIO-SPOTlight mouse enables a wide range of studies in all organ systems and functional monitoring opportunities not previously accessible.

## INTRODUCTION

The maintenance of cellular proteostasis is a dynamic and continual process achieved through regulation of multiple processes including protein synthesis and degradation, gene expression, and autophagy. The integrated stress response (ISR) pathway is a major signaling nexus for proteostatic maintenance under steady-state conditions and during exposure to a variety of environmental stressors. Four dedicated kinases (GCN2, HRI, PERK, and PKR) sense diverse forms of cellular stress including protein misfolding in the endoplasmic reticulum (ER), metabolic stress, oxidative stress, and viral infection (Pakos-Zebrucka et al., 2016). Each kinase phosphorylates the same serine residue on the eukaryotic translation initiation factor, eIF2α (*EIF2S1*). EIF2α phosphorylation functionally modifies protein translation by changing the delivery rate of methionine-charged initiator tRNA to scanning ribosomes. As the levels of p-eIF2α rise, general protein translation wanes but mRNAs containing certain regulatory upstream open reading frames (uORFs) are either resistant to this dip in translation or experience enhanced translation of the main, protein coding, ORF as part of ISR pathway activation. Enhanced translation of ISR effector proteins leads to downstream events that mount an adaptive response to stress, restoring cellular homeostasis or initiating apoptosis.

The core components of the ISR are highly conserved and have been subsequently found to support additional cellular processes not generally associated with cell stress. Regulating p-eIF2α levels and translation of one of its major effectors, the transcription factor ATF4, impacts multiple processes, including cell cycle progression, stem cell quiescence and pluripotency, and neurogenesis and axon guidance (Amiri et al., 2024; Cagnetta et al., 2019; Frank et al., 2010; Friend et al., 2015; Zismanov et al., 2016). Furthermore, the ISR pathway modifies synaptic plasticity associated with learning and memory as well as neuromodulation and the maintenance of circadian rhythms in the brain (Calakos & Caffall, 2024; Pathak et al., 2019; Sossin & Costa-Mattioli, 2019; Trinh & Klann, 2013). The ISR signaling network is also of great therapeutic interest as dysregulation of the ISR is implicated in both neurodevelopmental and neurodegenerative disorders and acts as a driving force in the progression of cancer (Lines et al., 2023; Lockshin & Calakos, 2024; Moon et al., 2018).Collectively, the above factors demonstrate the vital need for a detailed understanding of ISR pathway activity and how this translates to overall organismal health.

To date, understanding when and where ISR signaling occurs at the cellular level has faced technical challenges. Visually capturing cells with active ISR engagement has been difficult due to factors such as transitory phosphorylation states, low expression levels and rapid degradation of ISR effector proteins, and relatively few reliable commercial antibodies. Moreover, in some circumstances, the molecular hallmarks and functional effects of the ISR may dissociate (Guan et al., 2017), emphasizing the importance of state characterization at multiple levels. For these reasons, a variety of ISR reporters have been developed for use in cell culture and animal models (Chaveroux et al., 2015; Helseth et al., 2021; Iwawaki et al., 2017; Kang et al., 2015; Lu et al., 2004; Starck et al., 2016; Vasudevan et al., 2020; Vattem & Wek, 2004; Walter et al., 2015; Zhou et al., 2018). However, none of these reporters are available in a format for cell-type specific expression in a transgenic mouse line.

We previously developed the **S**elective **P**-eIF2α **O**RF **T**racking light, SPOTlight, reporter which replaces the ATF4 mRNA’s main ORF (mORF) and repressive uORF2 with cDNAs for fluorescent proteins (Helseth et al., 2021). The SPOTlight reporter offers multiple advantages and provides a distinctive readout of ISR-regulated function compared to molecular hallmarks of pathway activation. This is critical since adaptations as observed in chronic stress might dissociate these features (Guan et al., 2017). The SPOTlight readout directly reports the core functional consequence of ISR activation on translation initiation dynamics, and the ratiometric analysis of ORF translation (Red:Green fluorescence) reduces inherent variability associated with single-color reporters that could reflect either biological or technical variables.

Here, we expand on the capabilities of SPOTlight by developing it in a Cre-dependent format utilizing Double-floxed Inverted Orientation (DIO) of loxP recombination sites for cell-type specific expression and creating a transgenic mouse line. Through creation of a Cre-dependent targeted transgenic line, three major advantages are obtained – (1) the ability to selectively view distinct cell types over broad physical space, unrestricted by the limited spread of a viral vector, (2) the ability to distinguish activity in sparse and/or morphologically complex cells from surrounding tissue signal and (3) elimination of signal intensity variation inherent to viral delivery of genetic tools, related to targeting, tropism, and multiplicity of infection. Lastly, the use of a strong ubiquitous promoter (CAG) ensures high transcriptional abundance of reporter mRNA across most cell types, precluding the limitation of reporters that require endogenous *Atf4* mRNA to drive transcriptional readouts such as in the CARE-LUC model (Chaveroux et al., 2015).

This new resource can serve diverse investigational goals, such as quantitative comparison of specific cell populations distributed across broad spatial regions, discovery of novel cell subsets and instances of protein synthesis modulation, and for the first time, provides access to assessments that require identifying ISR pathway activity in living cells, such as measuring electrophysiological properties. In the studies that follow, we highlight these opportunities by providing example applications related to our interests in understanding the ISR in the central nervous system (CNS). We wish to note that many compelling applications are equally enabled by the DIO-SPOTlight mouse to explore other organ systems of particular relevance for the ISR, such as lung, pancreas, and liver.

## RESULTS

### Generation of DIO-SPOTlight transgenic reporter mice

The SPOTlight reporter design is based upon the 5’UTR of ATF4 mRNA, whose regulation by the ISR is extensively characterized (Dever et al., 2023; Lu et al., 2004; Pakos-Zebrucka et al., 2016; Vattem & Wek, 2004). The ATF4 5’UTR contains multiple uORFs: the very short uORF1 is self-contained upstream, whereas uORF2 overlaps out-of-frame with the ORF encoding the ATF4 protein and represses its translation when eIF2α is not phosphorylated. When the ISR is activated by eIF2α phosphorylation, the delivery rate of initiator methionyl tRNA to scanning ribosomes decreases. When this occurs, evidence supports that scanning ribosomes translate uORF1, then pass through uORF2’s start codon before receiving another initiator tRNA, resulting in less uORF2 translation inhibiting initiation at the main ATF4 coding ORF. In SPOTlight, enhanced green fluorescent protein (eGFP) cDNA is inserted after the uORF2 methionine start codon and tdTomato cDNA is inserted in place of the ATF4 coding sequence (***Figure 1A***).

**Figure 1.**
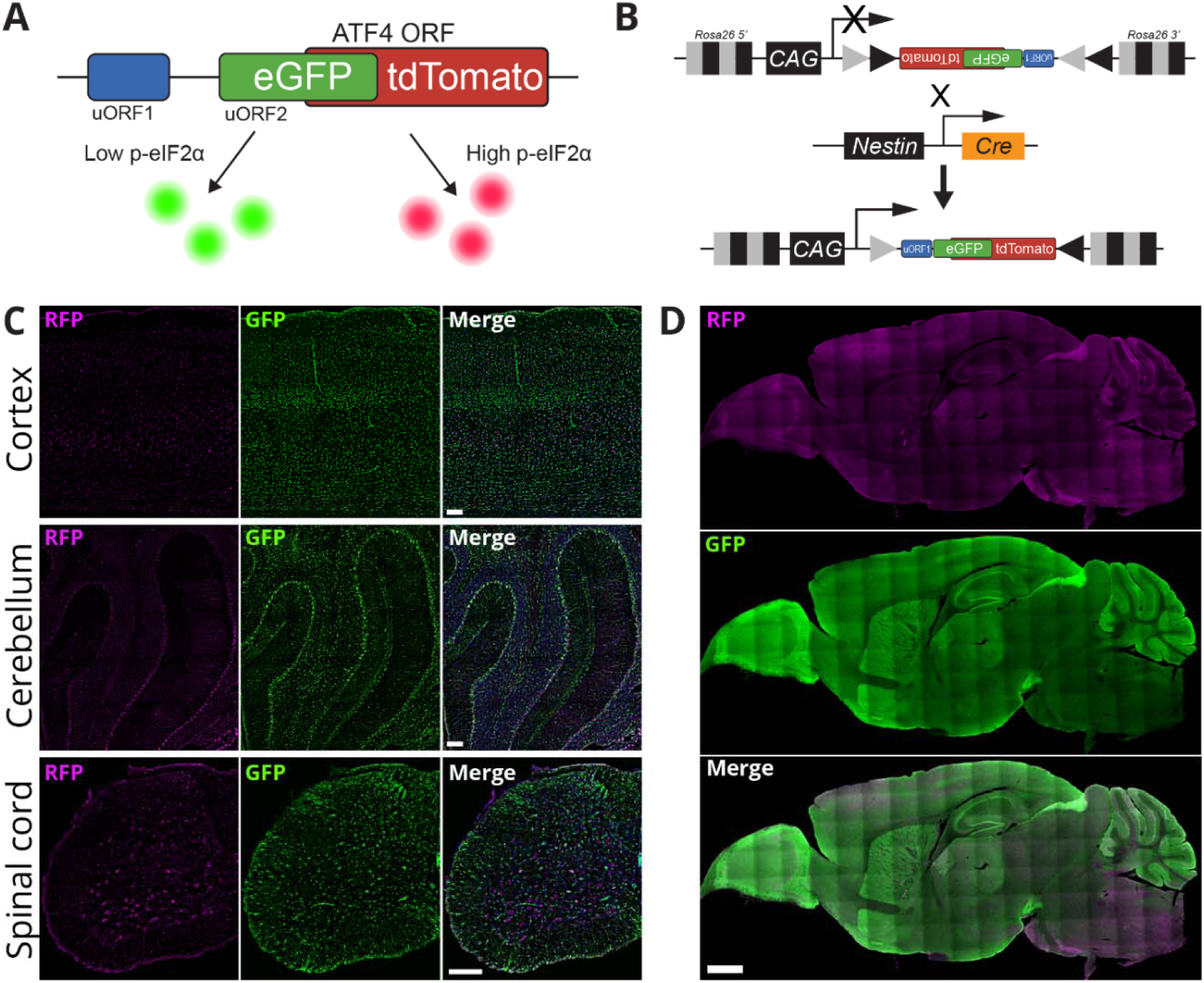
R26-DIO-SPOTlight design and expression in Nestin-Cre line. (**A**) SPOTlight reporter design. (**B**) R26-DIO-SPOTlight transgene insertion strategy and Nestin-Cre::DIO-SPOTlight mouse line generation. (**C & D**) Example images of expression in P30 mice. (**C**) Cortex (CTX), scale bar 100 µm. Cerebellum (CBM), scale bar 100 µm. Cervical spinal cord (SC), scale bar 200 µm. (**D**) Full sagittal brain slice, scale bar 1000 µm.

To generate DIO-SPOTlight, we modified SPOTlight to allow for Cre-dependent expression using a LoxP DIO system (Sohal et al., 2009). This design reduces the leaky expression seen using a traditional lox-Stop-lox cassette (Smedemark-Margulies & Trapani, 2013). In addition to the transgenic mouse line, DIO-SPOTlight cassette was cloned into an AAV packaging vector available from Addgene (submission in process to Addgene) (***Figure 1 – Supplement 1A-B***).

Generation of the Rosa26-DIO-SPOTlight transgenic mouse line (Figure 1B) was guided by the targeting strategy used for the Ai9 mouse line (Madisen et al., 2010). The R26Sor5’-pCAG-DIO-SPOTlight-WPRE-bGHpA-AttB-NeoR-AttP-R26Sor3’ cassette was targeted to the Rosa26 locus in hybrid 129S6/C57BL/6N (G4) embryonic stem cells and inserted through homologous recombination. Correctly targeted clones were selected by neomycin resistance and microinjected into morulae from the Institute of Cancer Research (ICR) mouse strain to produce chimeric mice. Chimeric founder mice were identified using PCR and DNA sequencing. Founder colonies were expanded then crossed with Nestin-Cre mice (Tronche et al., 1999) to achieve pan-neuronal expression throughout the CNS (***Figure 1B***). Nestin-Cre::DIO-SPOTlight hemizygous mice showed broad expression in the brain and spinal cord consistent with the expected pan-neuronal distribution of Nestin-Cre (***Figure 1C-D***). In general, green fluorescence was the dominant signal in cells, consistent with the understanding that the ISR is not activated in most cells of healthy normal tissue under basal conditions.

### Nestin-Cre::DIO-SPOTlight mice functionally differentiate rare subsets of cells

While qualitative visual inspection of brains from 30-day-old (P30) Nestin-Cre::DIO-SPOTlight mice showed a majority of cells with green signal predominating, there were also rare distinctive cells that were strongly red fluorescent without substantial green fluorescence (“red hot” cells, ***Figure 2A-C***). In many cases, these cells were distinguished by their unique SPOTlight readout from the majority of cells of similar cell type surrounding them. Importantly, these cells could also be identified in live tissue without fixation (***Figure 2A***).

**Figure 2.**
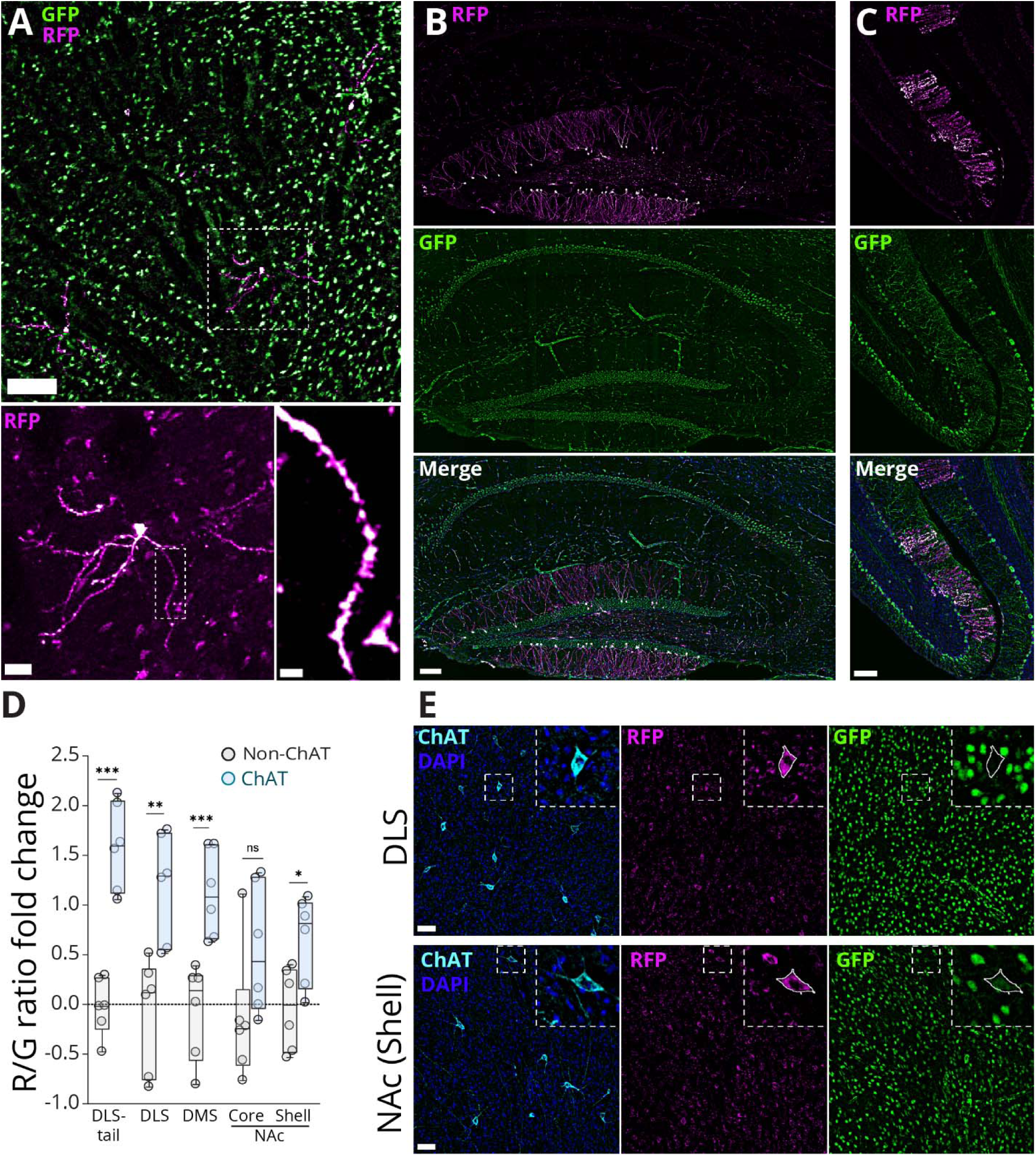
Nestin-Cre::DIO-SPOTlight fluorescence functionally differentiates rare subsets of cells. (**A-C**) Examples of “red hot” (strongly red, minimal green) neurons. Additional images available in ***Figure 2 – Supplement 1***. (**A**) Image from striatum in live acute brain slice tissue using multi-photon microscopy, Scale bars 100 µm, 20 µm, 5 µm. Remainder of images are of fixed tissue with immunohistochemistry (**B**, **C**, **E**). (**B**) Hippocampus, scale bar 100 µm. (**C**) Cerebellum, scale bar 100 µm. (**D**) Sub-regional SPOTlight red:green ratio analysis shown as log_2_ fold change of ChAT over non-ChAT neurons. N = 6 mice per region, DLS-Tail: ChAT(+) n = 41 cells, non-ChAT n = 128 cells; DLS: ChAT(+) n = 143 cells, non-ChAT n = 507 cells; DMS: ChAT(+) n = 154 cells, non-ChAT n = 519 cells; NAc core: ChAT(+) n = 93 cells, non-ChAT n = 434 cells; NAc shell: ChAT(+) n = 112 cells, non-ChAT n = 342 cells. Statistics summary tables and relative intensities of fluorescent proteins for panel **D** available in ***Figure 2 –Supplement 2***. (**E**) Representative images of dorsolateral striatum and nucleus accumbens shell, scale bars 50 µm.

In the dentate gyrus of hippocampus, visual scanning revealed that there were rare, dispersed cells distinguished by strong red SPOTlight signal but with morphology that was otherwise similar to neighboring granule cells. We show one striking example of a cluster of these cells (***Figure 2B***) providing an example of the utility of DIO-SPOTlight mice to highlight novel biology for follow-on studies. The dentate gyrus is unusual in that it is also a specialized area for neurogenesis. We wonder if these cells might share a “birthday” and/or might reflect a specific cell cycle state.

In cerebellum, we occasionally saw another unusual cluster of red-predominant cells, with distinctive morphology consistent with that of Bergmann glia (***Figure 2C***). This observation provides functional data to support a recent study that identified Bergmann glial cells with higher than normal ISR activation using gene expression profile data (Abbink et al., 2019). In addition, similar to what was previously observed with the viral reporter (Helseth et al., 2021), Nestin-Cre::DIO-SPOTlight mice revealed functional state variation across Purkinje cells.

Lastly, we recently reported that dorsal striatal cholinergic interneurons (CIN) have distinctive molecular hallmarks and functional SPOTlight activity indicative of steady-state ISR activity (Helseth et al., 2021). Here, we examined whether Nestin-Cre::DIO-SPOTlight mice reproduce those prior SPOTlight viral reporter observations and extended this analysis to consider whether this feature was present in other striatal subdivisions. Dorsal striatal subdivisions (medial, lateral, and tail) and ventral striatum (nucleus accumbens core and shell regions) each have distinctive input and output circuitry and functions. Using P30 Nestin Cre:DIO-SPOTlight mice, red and green fluorescence was measured in somata of cholinergic interneurons (ChAT+) and nearby non-ChAT+ cells in each striatal subdivision. Values obtained from Nestin-Cre::DIO-SPOTlight mice confirmed prior findings that dorsal striatal cholinergic interneurons show SPOTlight readouts characteristic of elevated ISR activity in comparison to neighboring non-ChAT neurons (Figure 2D-E). The mean ratio of red:green (R:G) fluorescence in dorsal striatal ChAT+ cells was more than twice the value of nearby non-ChAT cells (ChAT: 0.18 +/- 0.0074 SEM, n = 338 cells/6 mice, non-ChAT: 0.078 +/- 0.0082 SEM, n = 1154 cells/6 mice, P = 0.0001) yielding a log_2_ fold change of 1.221 +/- 0.18 SEM (P = 0.0001). Second, we found that this property was similar across striatal subdivisions, with all subdivisions except for the nucleus accumbens core showing statistically significant increases in R:G ratios for ChAT interneurons relative to neighboring non-ChAT neurons (***Figure 2 – Supplement 2***). These data are shown normatively using log2 fold change of the R:G ratio in ChAT+ cells relative to non-ChAT neighboring cells (***Figure 2D*** and ***Figure 2 – Supplement 2***) (DLS-Tail: 1.588 +/- 0.18 SEM, ChAT(+) n = 41 cells/6 mice, non-ChAT n = 128 cells/ 6 mice, P = 0.0002; DLS: 1.187 +/- 0.22 SEM, ChAT(+) n = 143 cells/ 6 mice, non-ChAT n = 507 cells/6 mice, P = 0.0072; DMS: 1.112 +/- 0.18 SEM, ChAT(+) n = 154 cells/6 mice, non-ChAT n = 519 cells/6 mice, P = 0.001; NAc core: 0.548 +/- 0.43 SEM, ChAT(+) n = 93 cells/6 mice, non-ChAT n = 434 cells/6 mice, P= 0.172; NAc shell 0.655 +/- 0.18 SEM, ChAT(+) n = 112 cells/6 mice, non-ChAT n = 342 cells/6 mice, P = 0.019). Finally, a multiple comparison analysis showed no significant differences in R:G ratios or normalized fold change between the ChAT populations in each subdivision. Thus, SPOTlight reporter values indicate that ISR-regulated protein synthesis initiation dynamics are increased throughout the striatum, while trends suggest there might be modestly higher activity in dorsal versus ventral subdivisions.

In summary, these examples demonstrate the utility of DIO-SPOTlight mice to identify distinctive functional properties in rare, distributed cell populations – whether these sparse cells are unanticipated and brought to attention by their SPOTlight “red hot” signal or are sparse and of specific interest, like striatal cholinergic interneurons which make up only 5% of striatal neurons.

### Acute in vivo tunicamycin exposure modifies SPOTlight readouts

We next determined the performance of Nestin-Cre::DIO-SPOTlight mice in the setting of an acute stressor that activates the ISR. Tunicamycin (3 µg/g), a drug that inhibits N-linked glycosylation of proteins and is a potent trigger of ER stress, or vehicle was delivered to P12 mice in 2 subcutaneous injections that were 2 hours apart, followed by euthanasia and brain tissue fixation 16-18h later (***Figure 3A***). In this experimental design, the R:G ratio was not significantly different between conditions, but did trend higher in the tunicamycin group (vehicle: 0.042 +/- 0.0015 SEM, n = 799 cells/5 mice; tunicamycin: 0.055 +/- 0.0016 SEM, n = 770 cells/5 mice, P = 0.145, ***Figure 3B***). We next examined the red and green components individually to provide further insight to their response kinetics under these conditions. Consistent with activation of the ISR, SPOTlight red fluorescence showed a highly significant increase to levels approximately 60% above vehicle (vehicle: 310 +/- 7.717 SEM, n = 799 cells/5 mice; tunicamycin: 505.6 +/- 9.297 SEM, n = 770 cells/5 mice, P = 0.0066, ***Figure 3C***). There was no significant difference in green fluorescence, but there was a slight trend towards increased values with tunicamycin treatment (vehicle: 8483 +/- 135.9 SEM, n = 799 cells/5 mice; tunicamycin: 11113 +/- 176.25 SEM, n = 770 cells/5 mice here, P = 0.1218, ***Figure 3D***). Representative images, including a SPOTlight transgene-negative mouse, are provided in ***Figure 3E***.

**Figure 3.**
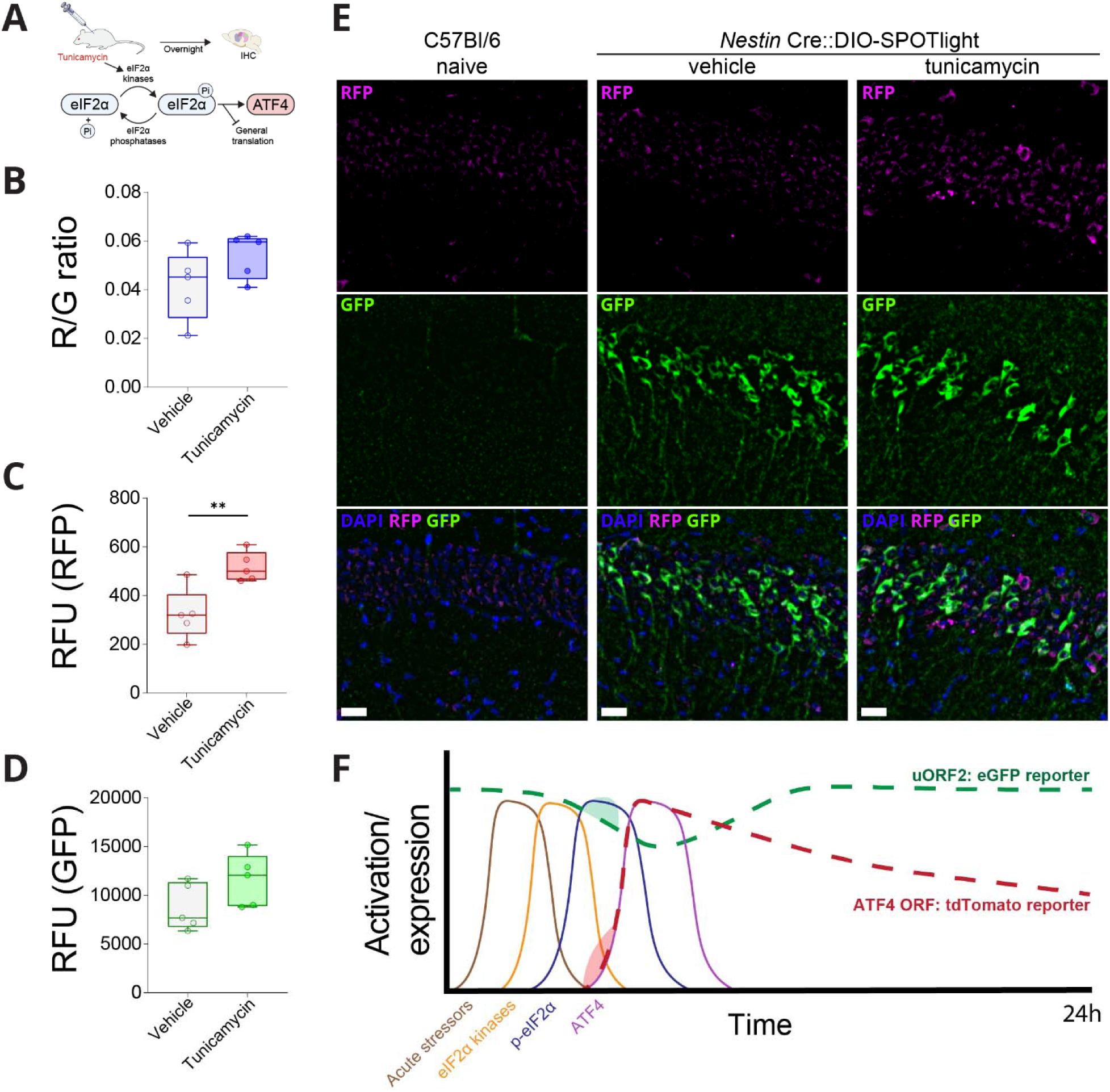
Acute tunicamycin exposure modifies SPOTlight readouts. (**A**) Schematic of ISR manipulation strategy. (**B-D**) Quantification of SPOTlight readouts: TM n = 5 mice, n = 771 cells; Vehicle n = 5 mice, n = 798 cells. (**B**) Red/green ratio: *t* = 1.614, *F*_(1,8)_ = 2.604, P = 0.1453. (**C**) Anti-RFP quantification *t* = 3.639, *F*_(1,8)_ = 13.24, P = 0.0066. (**D**) Anti-GFP quantification: *t* = 1.73, *F*_(1,8)_ = 2.994, P = 0.1218. (**E**) Representative images of red and green fluorescence in hippocampal pyramidal cell layer from transgene-negative control mice and Nestin Cre::DIO-SPOTlight hemizygous mice under vehicle and tunicamycin conditions. (**F**) Time course model for kinetics of ISR and SPOTlight reporter components after acute stressor/ISR activator. Following stressor, serial waves of eIF2α kinase activation, eIF2α phosphorylation, and ATF4 synthesis ensue. (Not shown, increased synthesis of other ISR effectors, phosphatases and downregulation of general protein synthesis.) Time course of SPOTlight EGFP translation is predicted to follow the period of general translation inhibition. Time course of SPOTlight tdTomato translation is predicted to follow ATF4 translation. Relative to the kinetics of deactivation and/or degradation of each ISR component, the half-lives of the SPOTlight reporters (tdTomato and EGFP) are relatively long (t_1/2_ approx. 24 h). In the particular case of tunicamycin as a stressor, there are predicted to be additional transient effects of inhibiting proteasomal degradation (Menéndez-Benito et al., 2005; Shenkman et al., 2007). The light shaded areas on each reporter are intended to indicate this effect which would cause additional accumulation of existing fluorophores relative to steady-state.

In summary, the increases in R:G ratio and red fluorescence mean values are consistent with predicted effects of ISR activation. However, the trend of an increase in mean green fluorescence was opposite to the predicted response to ISR activation. Although we waited 16 – 18 h from stressor to evaluation timepoint, the time course of this tunicamycin experiment is relatively short in comparison to the half-lives for degradation of existing fluorophores (i.e. eGFP, t ½ approx. 24h). We can therefore expect that some pre-existing eGFP that has not yet been degraded contributes to the R:G ratio. Perhaps more importantly, tunicamycin is known to also activate the unfolded protein response (UPR). Prior studies have shown that tunicamycin induction of the UPR causes a transient inhibition of the ubiquitin-protease system (UPS) and increased accumulation of cytosolic and ER proteins (Menéndez-Benito et al., 2005; Shenkman et al., 2007). This acute effect might relate to the trend for increased eGFP observed. This acute experiment provides an example of how SPOTlight components and ratio can vary according to the nature and acuity of an intervention, where varying contributions of proteins synthesized earlier may not yet be degraded (***Figure 3F***).

### CNS-wide Atlas of SPOTlight Activity in ChAT-positive Neurons

Lastly, we wanted to provide an example of using Nestin-Cre::DIO-SPOTlight mice to comprehensively survey cell types across distant regions and with heterogenous local tissue properties. We returned to the consideration of cholinergic neurons, now including populations of forebrain, cranial nerve and spinal cord nuclei. In comparing across more diverse tissue than intra-striatal regions, we sought to control for variations in tissue autofluorescence/background and SPOTlight signal related to surrounding neuropil density by using non-ChAT neighboring cell reference measures. We found that even though there was some variation in non-ChAT neighbor means across the regions, the vast majority of non-ChAT cell populations show mean R:G ratios around 0.1, whereas many ChAT populations have values more than twice that (***Figure 4A***). To normalize for variations in the non-ChAT neighbor means, we present a summary graph in which we have ordered the regions according to their normalized fold increase values measured relative to their non-ChAT neighbors (***Figure 4B***, same order is used in ***Figure 4A*** for direct comparison). From this analysis, it becomes apparent that most cholinergic neuron populations throughout the CNS show significantly elevated R:G ratios compared to their non-ChAT neighbors.

**Figure 4.**
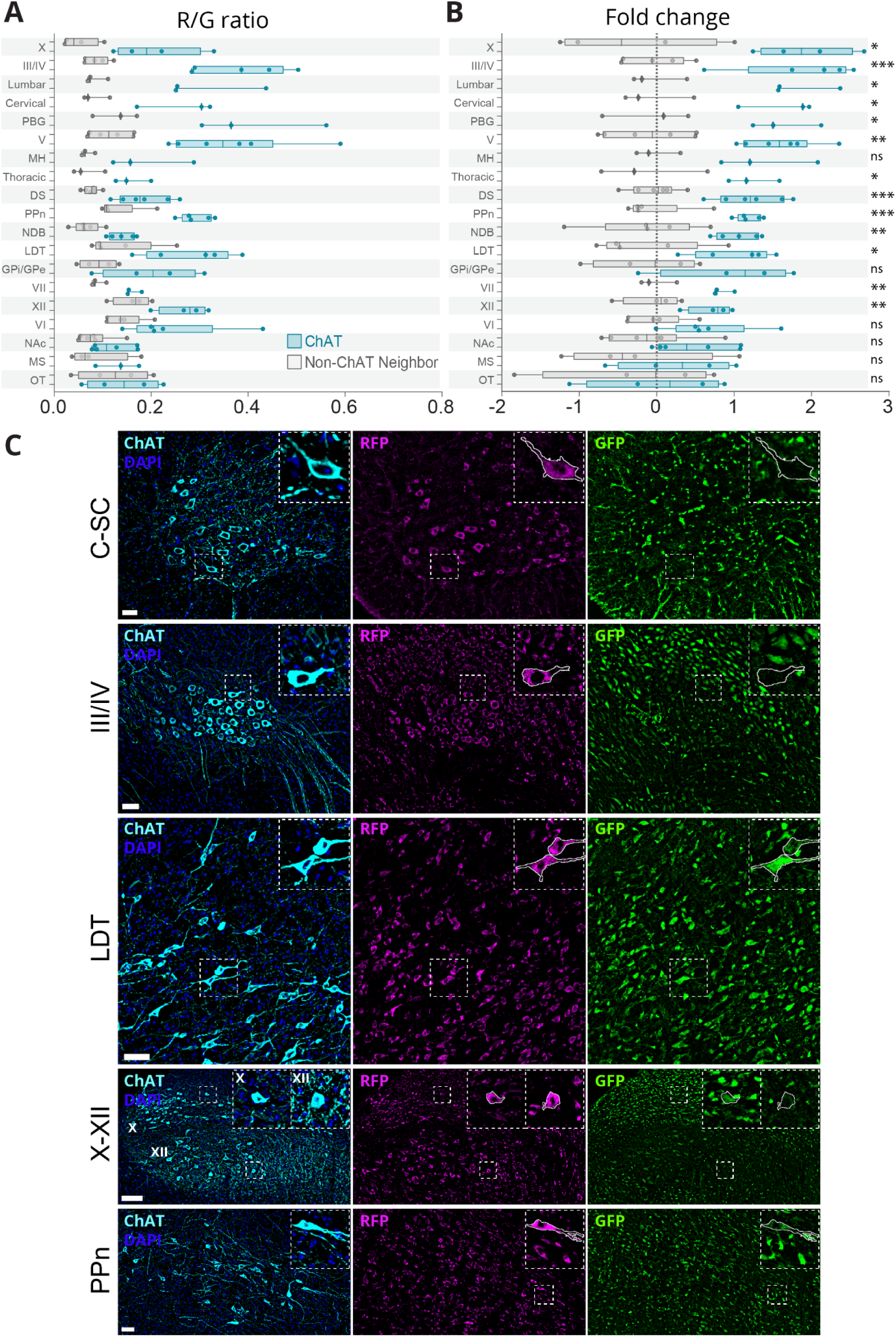
CNS-wide Atlas of SPOTlight Activity in ChAT-positive neurons. **(A)** Brain and spinal cord regional analysis of R:G ratio values in ChAT vs non-ChAT neighboring neurons. **(B)** Regional analysis of normalized Log2 (fold change) values of R:G ratios in ChAT(+)/non-ChAT neighboring neurons. **(A-B)** Statistical summary tables and relative intensities of fluorescent proteins for **A-B** available in ***Figure 4 – Supplement 1A-C***. *P < 0.05, **P < 0.01, ***P < 0.001, ****P < 0.0001. X: n = 4 mice, ChAT(+) n = 84 cells, Non-ChAT n = 66 cells; III/IV: n = 5 mice, ChAT(+) n = 144 cells, Non-ChAT n = 145 cells; Lumbar spinal cord n = 3 mice, ChAT(+) n = 250 cells, non-ChAT n = 165 cells; Cervical spinal cord n = 3 mice, ChAT(+) n = 257 cells, non-ChAT n = 182 cells; PBG: n = 3 mice, ChAT(+) n = 76 cells, Non-ChAT n = 69 cells; V: n = 6 mice, ChAT(+) n = 151 cells, Non-ChAT n = 136 cells; MH: n = 3 mice, ChAT(+) n = 241 cells, Non-ChAT n = 173 cells; Thoracic spinal cord n= 3 mice, ChAT(+) n = 342 cells, non-ChAT n = 210 cells; DS: n = 6 mice, ChAT(+) n = 338 cells, Non-ChAT n = 1154 cells; PPn: n = 5 mice, ChAT(+) n = 123 cells, Non-ChAT n = 200 cells; NDB: n = 5 mice, ChAT(+) n = 105 cells, Non-ChAT n = 144 cells; LDT: n = 5 mice, ChAT(+) n = 194 cells, Non-ChAT n = 241 cells; GPi/GPe: n = 4 mice, ChAT(+) n = 34 cells, Non-ChAT n = 77 cells; VII: n = 3 mice, ChAT(+) n = 91 cells, Non-ChAT n = 56 cells; XII: n = 4 mice, ChAT(+) n = 146 cells, Non-ChAT n = 101 cells; VI: n = 5 mice, ChAT(+) n = 52 cells, Non-ChAT n = 98 cells; NAc: n = 6 mice, ChAT(+) n = 205 cells, Non-ChAT n = 776 cells; MS: n = 4 mice, ChAT(+) n = 61 cells, Non-ChAT n = 111 cells; OT: n = 4 mice, ChAT(+) n = 39 cells, Non-ChAT n = 94 cells. **(C)** Representative images from select regions: Ventral horn of cervical spinal cord (VH), scale bar 50 µm; Oculomotor nucleus/trochlear nucleus (III/IV), scale bar 50 µm; Laterodorsal tegmental nucleus (LDT), scale bar 50 µm; Dorsal motor nucleus of the vagus nerve (X) and Hypoglossal nucleus (XII), scale bar 100 µm; Pedunculopontine nucleus (PPn), scale bar 50 µm. Example images for other regions are available in ***Figure 4 – Supplement 2***.

Further collapsing all cells measured brain-wide, group means illustrate that ChAT neurons have a significantly increased R:G ratio that is more than double that of non-ChAT neighbors (ChAT: 0.23 +/- 0.004 SEM, n = 2084 cells/6 mice; non-ChAT: 0.096 +/- 0.01 SEM, n = 3641 cells/6 mice, P = 0.0001). The ratio increase was accompanied by significant differences in both fluorophores, with ChAT+ neurons having significantly lower eGFP and higher tdTomato levels (tdT: ChAT: 1038 +/- 76.74 SEM, non-ChAT: 804 +/- 50.33 SEM, P = 0.03; EGFP: ChAT: 6073 +/- 196.6 SEM, non-ChAT: 10389 +/- 562.6 SEM, P < 0.0001). These SPOTlight reporter findings are consistent with the functional consequences predicted for ISR activation on *Atf4* mRNA reading frame dynamics (Dever et al., 2023). Finally, we did not observe any sex-dependent effects on SPOTlight measures, either in ChAT+ cells or non-ChAT cells (R:G ratios: ChAT male 0.24 +/- 0.045 SEM, n = 1155 cells/3 mice, female 0.23 +/- 0.005 SEM, n = 939 cells/3 mice, P = 0.437; Non-ChAT male 0.098 +/- 0.02 SEM, n = 1877cells/3 mice, female 0.094 +/- 0.013 SEM, n = 1764 cells/3 mice, P = 0.336, ***Figure 4 – Supplement 1A***).

## DISCUSSION

We have developed DIO-SPOTlight as a Cre-dependent reporter allowing for cell-type specific investigations of ISR-regulated protein synthesis dynamics. This new genetic tool is available in multiple formats, AAV vector and transgenic mouse, to allow flexibility for the specific needs of the researcher. In our assessments, we used DIO-SPOTlight mice crossed with Nestin-Cre to gain a broad neuronal view of SPOTlight activity and to perform a systematic quantitative comparison between ChAT and non-ChAT cell populations. In future applications, other experimental designs may restrict Cre expression to a single cell type and focus on questions of genotype, experience, exposure or developmental/age effects. In this study, we provided examples of the performance of DIO-SPOTlight measures in several application settings. These include visualization in unprocessed live tissue and quantitative comparisons of steady-state activity in a healthy tissue setting and in response to an acute cell stressor.

Historically, much that is known about the ISR’s function comes from immunohistochemical (IHC) readouts of molecular hallmarks of pathway activation, as well as a number of reporters (Helseth et al., 2021; Kang et al., 2015; Lu et al., 2004; Starck et al., 2016; Vasudevan et al., 2020; Vattem & Wek, 2004; Walter et al., 2015; Zhou et al., 2018) including two luciferase-based reporter mouse transgenic lines (Chaveroux et al., 2015; Iwawaki et al., 2017). These tools and techniques have provided indispensable understanding of this pathway. Prior to development of the DIO-SPOTlight mouse, investigators had access to a spatially coarse readout of luciferase bioluminescence in mouse lines that constitutively expressed either a reporter of ATF4 main ORF translation, UMAI mouse (Iwawaki et al., 2017) or ATF4 transcriptional activity, CARE-LUC mouse (Chaveroux et al., 2015).

The DIO-SPOTlight mouse and viral reporter resources presented here offer advantages that markedly advance ISR research techniques, and importantly, make it possible to identify specific cells with distinct states in live tissue for the first time. The ratiometric dual fluorophore readout of SPOTlight provides an internal control for reporter mRNA levels, overcoming a fundamental limitation inherent to single readout reporters. The Cre-dependent feature provides ready access to comprehensive examination of selected cell populations. Having the reporter available in a transgenic mouse line further markedly facilitates access to a number of broadly distributed, rare and/or morphologically complex cell types, e.g. microglia (which are also not amenable to most current viral transduction methods). In addition, there is less technical variation when using the mouse line because gene copy number is controlled for, in contrast to viral transduction and plasmid transfection methodologies.

ISR activity is often inferred through molecular hallmarks such as phosphorylation state of eIF2α or its kinases (e.g. PERK) or functional assays such as ATF4 transcriptional activity, global protein synthesis rates, and single ATF4 reading frame reporters. DIO-SPOTlight transgenic mice now provide an additional functional assay opportunity, with improved features that minimize technical variation and mechanistic ambiguity. A fair question is how to consider when one or the other approach may be more desirable. In living cells, the DIO-SPOTlight mouse provides an unparalleled opportunity to functionally visualize ISR activity. In fixed tissue settings, both IHC of molecular hallmarks of ISR activation and the SPOTlight readout are feasible and should be considered in conjunction. In IHC conditions, SPOTlight is particularly good for visual scanning to identify “hotspot” areas of activity variance. In contrast, the visual appearance of p-eIF2α IHC is more subtle, with diffuse punctate signal coming from surrounding neuropil difficult to distinguish from somata signal, making this a slower analytical process, that is not accessible to visual inspection alone.

Another important consideration is that the time course of each molecular hallmark and functional consequence of ISR activation has distinct and transient dynamics (***Figure 3F***). In this case, the long half-lives of eGFP and tdTomato (24h) serve to temporally extend readouts of ISR activity and can help identify cell populations that recently transiently engaged the ISR. However, a caveat of that extended time course is that manipulations on very acute time frames will include substantial amount of fluorophores that have not yet been degraded. Recognizing the differences in kinetics of each readout should be adequate for interpretation, as well as design, of experiments.

Besides a consideration of time course differences between ISR readouts, there may also be certain settings in which there is a dissociation between molecular hallmarks and functional readouts. One such example has been described under chronic ER stress, where p-eIF2α levels remained high, but functional readouts of protein synthesis were indicative of normal (non-ISR) conditions (Guan et al., 2017). In this example, compensation of eIF2 function by eIF3D was identified as the mechanism. These considerations highlight the value of a complementary suite of molecular and functional cell assessments.

The DIO-SPOTlight mouse line allowed us to follow up on previous findings that striatal cholinergic neurons exhibited elevated steady-state levels of ISR activation in the normal brain (Helseth et al., 2021). Using Nestin-Cre::DIO-SPOTlight mice, we replicated previous findings that dorsal striatal cholinergic neurons had elevated steady-state ISR engagement (***Figure 2D***) and extended this study to examine how broadly this feature was shared among other cholinergic neuron populations. Overall, we were surprised to see that this feature appeared to be widespread – including cholinergic interneurons and cholinergic projection neurons and diverse regions of basal forebrain, brain stem and spinal cord. When considered in aggregate, brain and spinal cord cholinergic neurons had significantly increased R:G ratios and red fluorescence and significantly decreased green fluorescence compared to neighboring neurons.

These differences are all consistent with translational effects of ISR activity. While relative fluorescence units (RFU) were higher in the brain than in the spinal cord, it is notable that the R:G ratios were nearly identical (R:G ratio - brain cholinergics 0.23 +/- 0.004 SEM, n = 2084 cells, 6 mice; spinal cholinergics 0.24 +/- 0.0088 SEM, n = 849 cells, 3 mice). The similarity of the R:G ratio but not absolute values of individual red and green components across two different CNS regions demonstrates the benefits of having a ratiometric reporter to normalize data with differences in total relative fluorescence whether the reasons are biological or technical.

The findings from this CNS-wide atlas of cholinergic neurons leads to the question: why are cholinergic neurons as a whole operating under an elevated ISR tone compared to non-cholinergic neurons? One obvious commonality is the synthesis, degradation, and release of acetylcholine. It is interesting to note that molecular chaperones are proposed to be the rate limiting step in the production of acetylcholinesterase (AChE), the molecule responsible for breaking down acetylcholine (Rotundo, 2017). In neuronal culture models, experiments have shown that ER stress-induced UPR activation led to a 3-fold increase in AChE expression (Liu et al., 2021). Beyond acetylcholine, cholinergic neurons are neuromodulatory cells that have expansive axonal arbors, high tonic secretory demand and utilize “volume” transmission as opposed to the highly temporally and spatially restricted synaptic transmission of neurotransmitters like glutamate and gamma-aminobutryic acid. Outside of the brain, cells with pronounced secretory roles depend heavily on the ER, showing evidence for chronic activation of the UPR that is thought to facilitate this secretory demand (Calfon et al., 2002; Reimold et al., 2001). These and other considerations provide rationale that additional neuromodulatory neuron types may also show high steady-state ISR engagement (discussed in Calakos & Caffall, 2024). Our previous work established physiological and behavioral consequences of ISR manipulations in striatal cholinergic interneurons (Helseth et al., 2021). The DIO-SPOTlight approach provides an opportunity to survey other neuromodulatory neuron classes and establish whether the ISR is also a behaviorally important modifier in those cells.

Some of the most visually striking observational findings in this study have been the presence of stochastic “red hot” neurons with little to no EGFP expression. These cells were observed in both live and fixed tissue in low numbers, sometimes as isolated cells or in clusters depending on the brain region. In neurons, besides reflecting cell stress, a red cell might reflect a cell that has recently undergone induction of synaptic plasticity, since ISR activation is associated with certain forms of long-term synaptic plasticity (Di Prisco et al., 2014). We further noted that “red hot” neurons were often seen in regions known for neurogenesis, i.e. the dentate gyrus, or known to receive migrating neural progenitors from the subventricular zone (SVZ), e.g. the olfactory bulb, striatum, and cortex (Jurkowski et al., 2020). Indeed, ISR signaling plays an important role in stem cell fate determination, neurogenesis, and axon guidance (Amiri et al., 2024; Cagnetta et al., 2019; Frank et al., 2010; Tahmasebi et al., 2019). DIO-SPOTlight provides an opportunity to identify cells that might not otherwise be distinguishable from their neighbors and further interrogate the mechanistic significance of their distinctive translational states.

In summary, DIO-SPOTlight transgenic mice provide a distinctive new functional readout to investigate known ISR biology and also to discover unexpected roles for the ISR. For known ISR biology, DIO-SPOTlight transgenic mice are well-suited for quantitative comparisons because transgene copy number is controlled and the ratiometric readout provides an internal control for mRNA levels. Cre-dependent expression further allows for the visualization of selected subsets of cells in isolation from surrounding cellular signal. For novel discovery, we have already found a number of intriguing cell subsets that stand out because of their red predominance from this first look at young Nestin-Cre::DIO-SPOTlight mice. There remains a large number of applications for DIO-SPOTlight mice over developmental time, disease state, exposures, learning experiences and organ systems that should be further revealing. Taken together, our study demonstrates the effective use of a new resource, DIO-SPOTlight, and provides examples of how this tool may provide great insights into ISR engagement in a variety of contexts.

## MATERIALS AND METHODS

The data, code, protocols, and key lab materials used and generated in this study are listed in a Key Resource Table alongside their persistent identifiers at 10.5281/zenodo.16368495.

### Viral construct

The original SPOTlight construct was published in Helseth et al., 2021. Briefly, SPOTlight was an in-house designed reporter consisting of recombinant DNA based on the known published sequence of murine *Atf4* mRNA (NCBI Reference Sequence: NM_009716.3). To achieve Cre-recombinase dependent expression, SPOTlight was cloned into AAV vector driven by the CAG promoter in a double-floxed inverted orientation (DIO) containing two pairs of incompatible Lox sites (LoxP and Lox2722) by Vector Builder (https://en.vectorbuilder.com/).

### Animals

Rosa26-DIO-SPOTlight mouse was produced by first cloning DIO-SPOTlight construct into Ai9 vector backbone in place of the Lox-STOP-Lox-tdTomato cassette (Ai9 was a gift from Hongkui Zeng; Addgene plasmid # 22799; http://n2t.net/addgene:22799; RRID:Addgene_22799) by Vector Builder (https://en.vectorbuilder.com/). Linearized Ai9-DIO-SPOTlight plasmid was then transferred to Duke Transgenic Mouse Core (Rodent Genetic Engineering Services). Integration into the Rosa26 locus was achieved through homologous recombination following electroporation into hybrid 129S6/C57BL/6N (G4) embryonic stem cells. Pan-neuronal expression of DIO-SPOTlight was achieved by crossing Rosa26-DIO-SPOTlight hemizygous transgenic founders (Founder lines ID: C1 and D8) with Nestin-Cre mice, obtained from Jackson Laboratories; JAX: strain #003771, RRID: IMSR_JAX:003771 (Tronche et al 1999). Two founder lines were maintained and used in experiments as follows: brain-wide regional analysis of cholinergic neurons n = 6 (5 founder D8 and 1 founder C1, age P30 – P33), spinal cord cholinergic neuron analysis n = 3 (2 founder D8 and 1 founder C1, age P30), and in vivo tunicamycin injection n = 10 (tunicamycin – 5 founder C1, vehicle – 4 founder C1 and 1 founder D8, age P13). All experimental procedures were approved by the institutional animal care and use committee at Duke University. Animals were maintained on a 12:12 hour light:dark cycle with ad libitum access to food and water.

### Genotyping

Genotyping was performed using polymerase chain reaction of genomic DNA with the following primers: R26 wild type (WT) forward 5’-AAG GGA GCT GCA GTG GAG TA-3’; WT reverse 5’-CCG AAA ATC TGT GGG AAG TC-3’; DIO-SPOTlight forward 5’- AAC TGT GGC GTT AGA GAT CG-3’; DIO-SPOTlight reverse 5’-TTG AGC TCA GGA ACC TAT AAC T-3’. Cre genotyping was performed using the generic Cre protocol 22392 (The Jackson Laboratory, Bar Harbor, ME).

### In vivo tunicamycin treatment

Tunicamycin (TM) was purchased from Tocris (Bristol, UK) and dissolved in dimethyl sulfoxide (DMSO) to a concentration of 5 mg/ml. On the day of the experiment, it was further diluted 1:25 (v/v) in 150 mM dextrose in PBS with 5% Tween-80 to maintain TM solubility and administered at a final concentration 3 µg/g bodyweight per mouse. Following methodology reported in Wang et al 2015, twelve-day old mice received two subcutaneous injections spaced two hours apart beginning at 3:00 PM. The mice were sacrificed 16 – 18 hours after the final injection, and the dissected brains were stored for further processing. Imaging and data analysis was performed blind to treatment groups.

### Tissue processing

Mice were anesthetized by using a drop jar method containing 3-5% aerosolized isoflurane. The thoracic cavity was then opened and transcardial perfusion with 1x PBS followed by fixation with 4% paraformaldehyde (PFA) was performed. The perfused brains were then removed and stored in 4% PFA + 20% sucrose for 24 hours at 4LC. The brains were then transferred to 30% sucrose in PBS for an additional 24 hours before embedding in Tissue Tek optimal cutting temperature compound (OCT, Torrance, CA) and stored at -20LC until sliced into 40 μm sagittal slices using a Leica CM 1950 cryostat. Brain slices were stored in PBS with 0.01% sodium azide at 4LC.

After removing the brain, spinal cord dissection occurred using the following landmarks: cervical, base of skull to caudal T1 lamina; thoracic, 0.6 cm segment centered on T9 lamina; and lumbar, identified based on L1-L6 nerve roots. Spinal cord tissue was post-fixed using cyroPREP (Covaris, Woburn, MA) in 20% sucrose overnight, 30% sucrose for 5 days, then embedded in OCT, frozen on dry ice, and stored at –80°C. Tissue blocks acclimated at –20°C overnight prior to serial cryo-sectioning in the transverse plane to generate 20µm-thick cross sections using Microm HM 505N instrument.

### IHC

Brain and spinal cord IHC differed only in that the brain slices were floating, and spinal cord sections were pre-mounted on slides. Any remaining free aldehyde groups were quenched with 0.1M glycine in 1x PBS for 30 min at room temperature (RT). Alkaline retrieval was then performed for 20 minutes at RT using freshly prepared 1% NaOH in 1x PBS while gently shaking. The sections were then washed three times for 10 minutes in PBS-T (1x PBS with 0.3% Triton X-100) and blocked for one hour in 10% normal donkey serum (NDS) plus 1% bovine serum albumin (BSA) in PBS-T, all while gently shaking on a VWR Standard Analog Shaker on low speed at RT. Slices were then incubated in primary antibodies on a shaker at 4LC for 24 hours then washed three times in PBS-T at RT for 10 minutes each wash. Corresponding secondary antibodies were then added to the slices and incubated for 1 hours on a shaker at RT. Secondary antibodies were then removed and nuclei were then labeled for 10 minutes with Hoechst 33342 (Enzo Life Sciences) at a final concentration of 10 μg/mL in PBS-T. This was followed with two additional 10-minute washes in PBS-T at RT while on a shaker. PBS-T was exchanged for 1 x PBS and the slices were placed on glass slides and coverslips were sealed with Vectashield Vibrance Mounting Media (cat no. H-1700). All antibodies were prepared in dilution buffer containing 5% NDS plus 1% BSA in PBS-T. Commercially available antibodies used were anti-RFP (rabbit,1:1000, Rockland Antibodies and Assays, cat no. 600-401-379; RRID: AB_2209751), anti-GFP (chicken, 1:1000, Abcam, cat no. AB13970, RRID: AB_300798), and anti-ChAT (goat, 1:200, Millipore, cat no. AB144P; RRID: AB_2079751). Corresponding secondary antibody conjugates Alexa Fluor 488, Alexa Fluor 546, and Alexa Fluor 647 raised in donkey were obtained through Invitrogen (RRID: AB_2921070; cat no. A78948, RRID: AB_2534016; cat no. A10040, and RRID: AB_25335864; cat no. A32849, respectively)

### Confocal image acquisition and processing

Fixed tissue sections were imaged using a Leica DMi8 Andor Dragonfly 202 unit equipped with Borealis illumination spinning disk confocal microscope and iXon 888 EMCCD camera. The full brain image was captured using a 10x objective lens plus a 1.5x camera zoom. All other regions of interest (ROI) were imaged using a 40x/1.3 oil emersion objective lens. For ratiometric SPOTlight analysis laser power and exposure time settings were held constant across acquisition channels for tdTomato and EGFP. Completed multi-framed images were stitched together using Fusion Stitcher (Oxford Instruments). Post-acquisition micrographs were analyzed using FIJI ImageJ analysis software (http://imagej.nih.gov/).

For measures of cholinergic and non-cholinergic neighboring neurons, 19 regions were imaged in P30 mice tissue. For acute tunicamycin experiments, methods are the same as described for the cholinergic and non-cholinergic analysis, with the exception that hippocampal pyramidal neurons from P13 mice were analyzed. Cholinergic neurons were used to create a threshold intensity cell body mask from a single z-plane using anti-ChAT as a cellular marker. ChAT negative neurons were selected using the SPOTlight reporter as a cell body mask. Given the two-fluorophore nature of the SPOTlight reporter and to limit biasing cell identification based on signal in one channel over the other, the image pixel values from the 561nm and 488nm channels were added together using FIJI’s image calculator option. Background subtraction for cell masking was performed by applying a 15-pixel gaussian blur on a duplicate image, subtracting the pixel values of blurred image from the original, and then applying a second 2-pixel gaussian blur on the resulting background subtracted image. Cell body ROIs were then selected using the wand selection tool and ROI manager window. In some cases, where cell bodies were clearly definable but high background prevented accurate thresholding, ROIs were drawn manually. Cytosolic signal analysis was achieved through nuclear exclusion in one of three ways. First, in many cases the nuclear compartment was clearly definable through the primary masking channel, selected, and added to the ROI manager. When this was not available, a nuclear mask was made from the Hoechst stain as described above for the primary masking channels. For both circumstances, the nuclear compartment was excluded from the analysis using the XOR function available in ROI manager’s “More” drop down window. The third method was used only when the primary cell body thresholding did not fully enclose the nuclear compartment within mask. The ROIs were then used to measure regions from the original image plane without need to exclude the nucleus since that region was not included in the ROI. Within slice background fluorescence was controlled for by measuring red and green fluorescence values in nearby cells where recombination did not occur (negligible red and green signal). These cells were identifiable in all regions, as Nestin-Cre recombinase efficiency was not 100%. Five such cells were selected for each region. The average red and average green signals from these SPOTlight-negative somata were subtracted from all cell measures.

### Live tissue preparation and two-photon imaging

To prepare acute brain slices, an adult animal (∼2 months old) was anesthetized and intracardially perfused with an ice-cold *N*-methyl-D-glucamine (NMDG)–based ACSF solution containing the following (in mM): 90 NMDG, 2.5 KCl, 1.2 NaH_2_PO_4_, 35 NaHCO_3_, 20 HEPES, 25 glucose, 5 sodium ascorbate, 2 thiourea, 3 sodium pyruvate, 10 MgSO_4_ heptahydrate, 0.5 CaCl_2_ dihydrate, and saturated with 95% O_2_ and 5% CO_2_. The brain was then quickly removed and 250-μm sagittal sections were cut on a Leica VT1200S vibratome. Immediately after cutting, slices were placed in a 32°C recovery chamber containing NMDG ACSF for 10 to 15 min. After heated recovery, live tissue brain slices were transferred to and maintained in a holding chamber containing artificial cerebrospinal fluid (ACSF; 124 mM NaCl, 2.5 mM KCl, 1 mM MgCl_2_, 2 mM CaCl_2_, 26 mM NaHCO_3_, 1.2 mM NaH_2_PO_4_, and 10 mM glucose, saturated with 95% O_2_ and 5% CO_2_, pH 7.4, 300 mOsm/l). Slices rested for approximately 1 hour prior to imaging.

For imaging, slices were placed in a perfusion chamber and superfused with oxygenated ACSF. Slices were imaged using a commercial two-photon microscope (Bruker Investigator Multiphoton Imaging System). Imaging of SPOTlight in live tissue slices was performed using 920nm light emitted from a Coherent Chameleon I Ti:Sapphire femtosecond laser with the laser power kept between 35-50 mW after the objective. Fluorescence signal was collected using a Nikon CFI75 LWD 16× W Objective and Quad GaAsP photomultiplier tubes (PMTs). Sample SPOTlight signal was split into red and green channels through a 565lpxr dichroic mirror and then passed through a 525/70 bandpass PMT filter. An image stack (512 x 512 resolution) of ∼200 µm total depth was collected using galvo-galvo scanning with frame rate of 1 Hz with 3-4 µm depth increments at either 1X or 4X optical zoom.

### Statistical analysis

All statistics were performed by nesting measured cells within the mouse they were collected from with a significance level set at *P* < 0.05 using the two-tailed nested *t* test option in GraphPad Prism 10.3 software (GraphPad Software, Boston, MA). P value summary asterisk values used were *P < 0.05, **P < 0.01, ***P < 0.001, ****P < 0.0001. Graphical depiction of the data is shown as averages from all cells per mouse in 5-95 percentile box and whisker plots overlayed with circular symbols indicating the mean value from each individual mouse. 4/1573 cells were excluded as outliers from the data for figure 3B-D and 1/7,132 cells was excluded as an outlier from the data for Figure 4. Outliers were defined as being +/- 5SD from the mean within mouse.

## Supporting information

Supplemental Data Files_DIOSPOTlight

## Acknowledgements

We thank Victoria Hall for preparing and obtaining live tissue images, Kaylee Bayles for mouse genotyping and mouse colony assistance, Briana Cellini for technical assistance with spinal cord processing, Jessica Matsuoka for technical assistance with spinal cord processing and methods, and Sharon Powley and Joseph Rittiner for critical reading of manuscript drafts.

## ADDITIONAL INFORMATION

### Competing interests **None.**

### Author Contributions

M.L.O. performed all SPOTlight imaging in Nestin-Cre::DIO-SPOTlight fixed tissue samples, designed experiments and injected mice for ISR manipulation, brain tissue preparation, designed immunohistochemical experiments and immunohistochemical staining of floating brain sections and data interpretation. Z.F.C. designed AAV-DIO-SPOTlight and the R26-DIO-SPOTlight mouse and aided in conceptual development, data interpretation, statistical analysis and graphical depiction of data. T.D.F. performed spinal cord dissection, tissue preparation, immunohistochemical staining of mounted spinal cord section, and provided expertise on spinal cord anatomy. C.B.E. performed early foundational experiments using AAV-DIO-SPOTlight to measure striatal cholinergic interneurons in striatal subregions and data for Figure 1 – Supplement 1. M.L.O. and N.C. wrote the manuscript with input from the other authors. N.C. provided conceptual development, research oversight, and data interpretation.

