## Supplemental Data Files_DIOSPOTlight for "DIO-SPOTlight Transgenic Mouse to Functionally Monitor Protein Synthesis Regulated by the Integrated Stress Response"

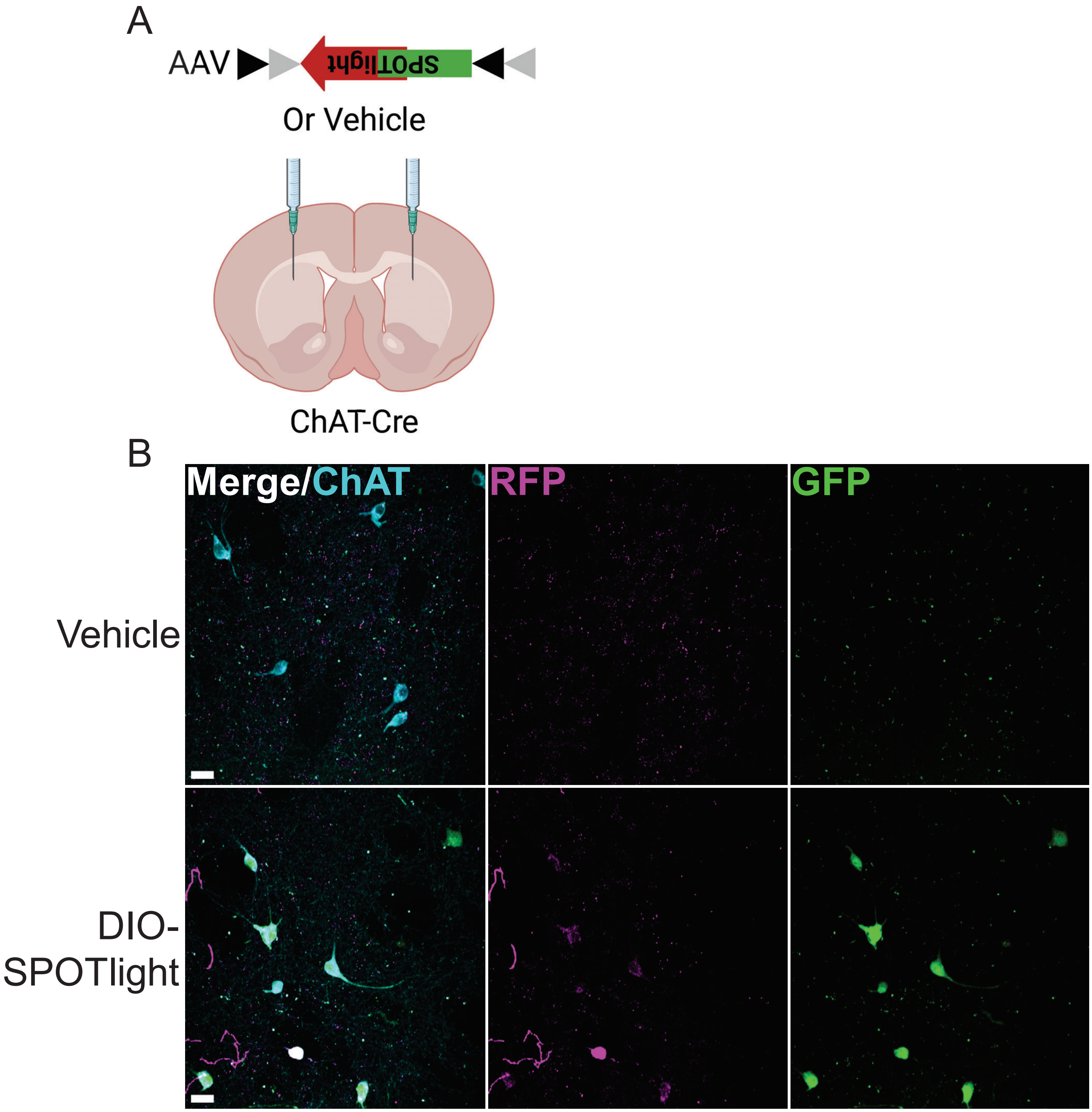


**Figure 1 — Supplement 1**. AAV-DIO-SPOTlight. (**A**) Injection strategy for Cre-dependent expression of AAV-DIO-SPOTlight in a ChAT-Cre mouse line (created in BioRender). (**B**) Representative images of AAV-DIO-SPOTlight expression and contralateral vehicle injection.

**Figure 2 — Supplement 1.** Additional examples of "red hot" neurons found in different brain regions. CTX – Cortex, scale bar 50 µm. NAc – Nucleus accumbens, scale bar 25 µm. DS – Dorsal striatum, scale bar 50 µm. OB – Olfactory bulb, scale bar 50 µm. HPC – Hippocampus, scale bar 50 µm.

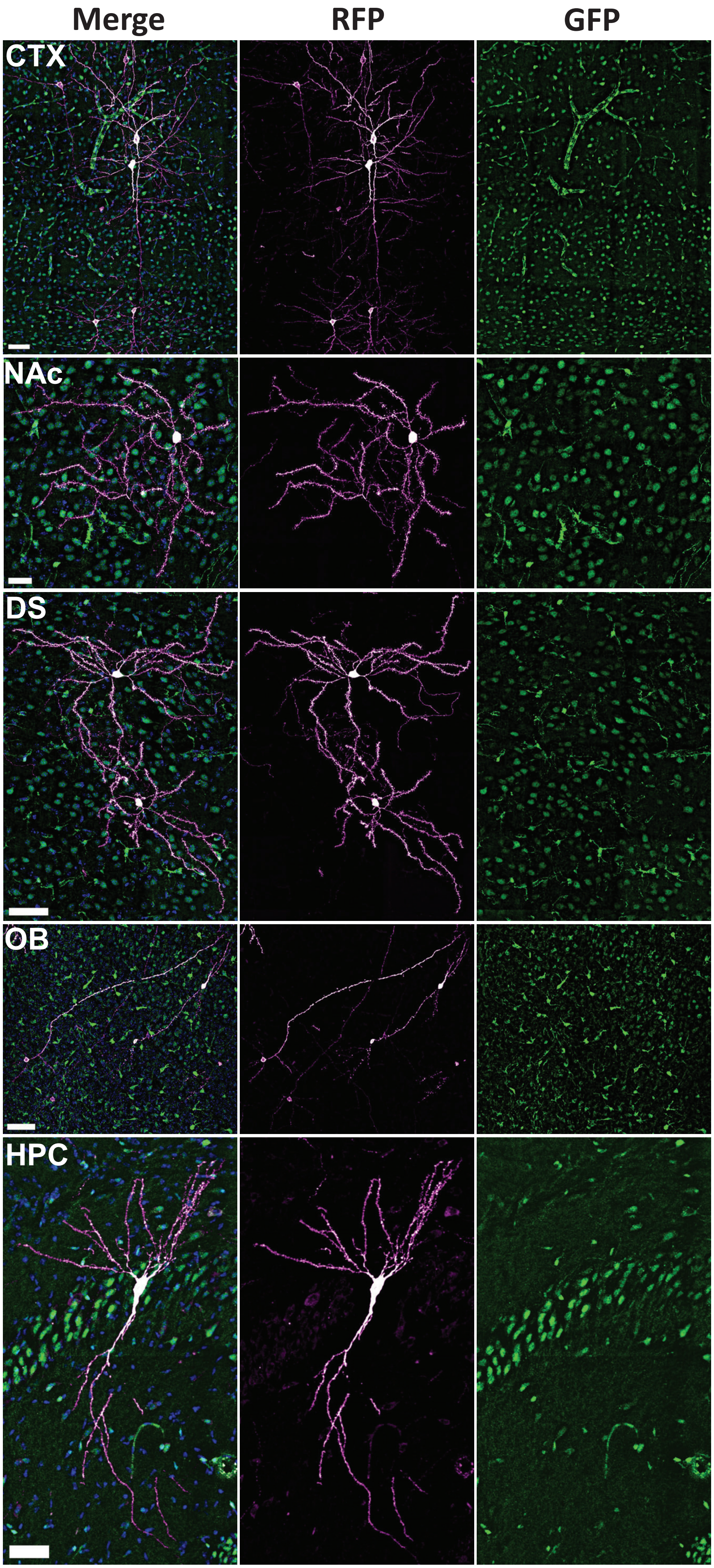

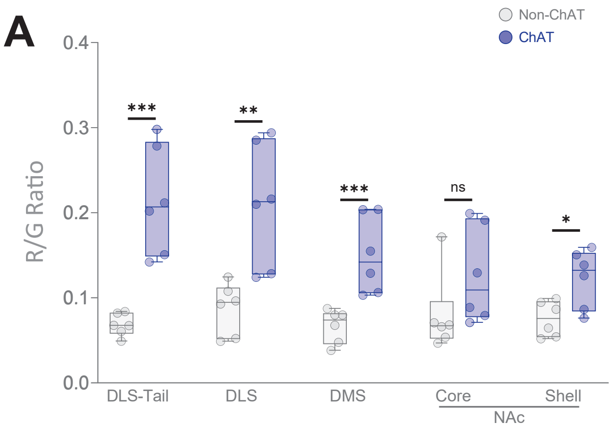


**Figure 2 Statistical Summary (Nested t test: R/G Ratios and $Log_2$ Fold Change)**

| Non-ChAT vs ChAT | P value | P value summary | P < 0.05? | One- or two-tailed P value? | t, df | F, DFn, Dfd | R:G Ratio mean difference ± SEM | $Log_2$ FC mean difference ± SEM |
|---|---|---|---|---|---|---|---|---|
| Str DLS tail | 0.0002 | *** | Yes | Two-tailed | t=5.600, df=10 | 31.35, 1, 10 | -0.1431 ± 0.02556 | 1.588 ± 0.179 |
| Str DMS | 0.001 | *** | Yes | Two-tailed | t=4.595, df=10 | 21.11, 1, 10 | -0.08107 ± 0.01764 | 1.112 ± 0.179 |
| Str DLS | 0.0072 | ** | Yes | Two-tailed | t=3.365, df=10 | 11.33, 1, 10 | -0.1131 ± 0.03361 | 1.187 ± 0.221 |
| NAc Shell | 0.0188 | * | Yes | Two-tailed | t=2.799, df=10 | 7.835, 1, 10 | -0.04662 ± 0.01665 | 0.655 ± 0.18 |
| NAc Core | 0.1721 | ns | No | Two-tailed | t=1.471, df=10 | 2.163, 1, 10 | -0.03940 ± 0.02679 | 0.548 ± 0.43 |

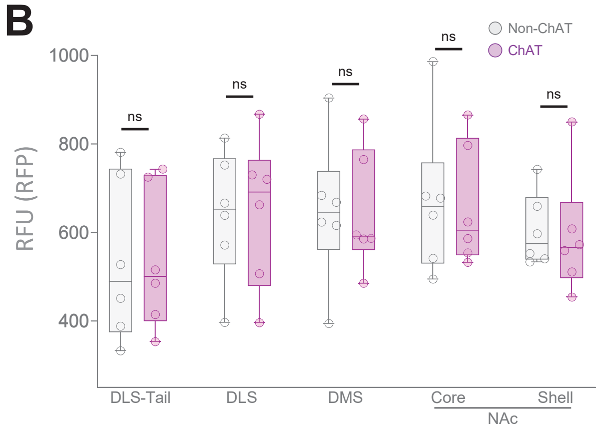


**Figure 2 Statistical Summary (Nested t Test: RFP Relative Abundance)**

| Non-ChAT vs ChAT | P value | P value summary | P < 0.05? | One- or two-tailed P value? | t, df | F, DFn, Dfd | Mean difference ± SEM |
|---|---|---|---|---|---|---|---|
| NAc Shell | 0.6985 | ns | No | Two-tailed | t=0.3986, df=10 | 0.1589, 1, 10 | 23.44 ± 58.79 |
| Str DLS | 0.8153 | ns | No | Two-tailed | t=0.2399, df=10 | 0.0576, 1, 10 | 26.57 ± 110.7 |
| NAc Core | 0.9001 | ns | No | Two-tailed | t=0.1287, df=10 | 0.0166, 1, 10 | 11.65 ± 90.51 |
| Str DMS | 0.9435 | ns | No | Two-tailed | t=0.0726 3, df=10 | 0.00538 , 1, 10 | -6.774 ± 93.27 |
| Str DLS tail | 0.9706 | ns | No | Two-tailed | t=0.0379, df=10 | 0.00143 , 1, 10 | -3.792 ± 100.2 |

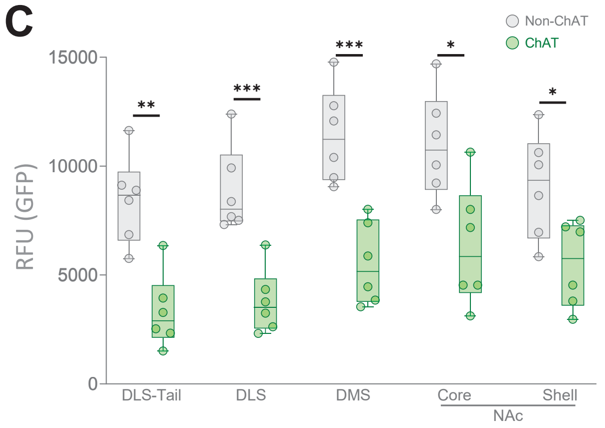


**Figure 2 Statistical Summary (Nested t Test: GFP Relative Abundance)**

| Non-ChAT vs ChAT | P value | P value summary | P < 0.05? | One- or two-tailed P value? | t, df | F, DFn, Dfd | Mean difference ± SEM |
|---|---|---|---|---|---|---|---|
| Str DLS | 0.0005 | *** | Yes | Two-tailed | t=5.120, df=10 | 26.22, 1, 10 | 5109 ± 997.8 |
| Str DMS | 0.0006 | *** | Yes | Two-tailed | t=4.957, df=10 | 24.58, 1, 10 | 5883 ± 1187 |
| Str DLS tail | 0.0011 | ** | Yes | Two-tailed | t=4.532, df=10 | 20.54, 1, 10 | 5070 ± 1119 |
| NAc Core | 0.0118 | * | Yes | Two-tailed | t=3.074, df=10 | 9.448, 1, 10 | 4603 ± 1497 |
| NAc Shell | 0.0175 | * | Yes | Two-tailed | t=2.842, df=10 | 8.076, 1, 10 | 3619 ± 1273 |

**Figure 2 — Supplement 2.** Sub-regional analysis of striatum. (**A-C**) N = 6 mice per region. DLS-tail ChAT: n = 41 cells, non-ChAT: n = 128 cells; DLS ChAT: n = 41 cells, non-ChAT: n = 128 cells; DMS ChAT: n = 154 cells, non-ChAT: n = 519 cells; NAc core ChAT: n = 93 cells, non-ChAT: n = 434 cells; NAc shell ChAT: n = 93 cells, non-ChAT: n = 434 cells. (**A**) R:G ratios with statistical table. The underlying data between R:G ratios and $log_2$ fold-change are the same with the same statistical summaries with the exception of mean differences. (**B**) Relative red fluorescence abundance with statistical table. (**C**) Relative green fluorescence abundance with statistical table.

**A**

**Figure 4 Statistical Summary (Nested t test: R/G Ratios and $Log_2$ Fold Change)**

| Non-ChAT vs ChAT | P value | P value summary | P < 0.05? | One- or two-tailed P value? | t, df | F, DFn, Dfd | R:G Ratio mean difference ± SEM | $Log_2$ FC mean difference ± SEM |
|---|---|---|---|---|---|---|---|---|
| DS | 0.0001 | *** | Yes | Two-tailed | t=5.919, df=10 | 35.03, 1, 10 | -0.1026 ± 0.01734 | 1.216 ± 0.178 |
| III/IV | 0.0002 | *** | Yes | Two-tailed | t=6.562, df=8 | 43.06, 1, 8 | -0.2945 ± 0.04488 | 1.8844 ± 0.315 |
| PPn | 0.0003 | *** | Yes | Two-tailed | t=6.192, df=8 | 38.34, 1, 8 | -0.1642 ± 0.02652 | 1.187 ± 0.067 |
| V | 0.0011 | ** | Yes | Two-tailed | t=4.536, df=10 | 20.57, 1, 10 | -0.2491 ± 0.05493 | 1.586 ± 0.199 |
| NDB | 0.0025 | ** | Yes | Two-tailed | t=4.346, df=8 | 18.89, 1, 8 | -0.07492 ± 0.01724 | 1.05 ± 0.16 |
| VII | 0.0045 | ** | Yes | Two-tailed | t=5.780, df=4 | 33.41, 1, 4 | -0.07318 ± 0.01266 | 0.8433 ± 0.057 |
| XII | 0.0092 | ** | Yes | Two-tailed | t=3.780, df=6 | 14.29, 1, 6 | -0.1227 ± 0.03247 | 0.7147 ± 0.121 |
| LDT | 0.0207 | * | Yes | Two-tailed | t=2.874, df=8 | 8.260, 1, 8 | -0.1489 ± 0.05181 | 1.016 ± 0.21 |
| Lumbar - no outlier | 0.0241 | * | Yes | Two-tailed | t=3.537, df=4 | 12.51, 1, 4 | -0.2272 ± 0.06422 | 1.84 ± 0.445 |
| X | 0.0262 | * | Yes | Two-tailed | t=2.933, df=6 | 8.605, 1, 6 | -0.1627 ± 0.05546 | 1.915 ± 0.253 |
| Cervical | 0.0264 | * | Yes | Two-tailed | t=3.434, df=4 | 11.79, 1, 4 | -0.1813 ± 0.05279 | 1.635 ± 0.418 |
| PBG | 0.0274 | * | Yes | Two-tailed | t=3.394, df=4 | 11.52, 1, 4 | -0.2818 ± 0.08302 | 1.622 ± 0.185 |
| Thoracic | 0.0348 | * | Yes | Two-tailed | t=3.142, df=4 | 9.874, 1, 4 | -0.09142 ± 0.02909 | 1.223 ± 0.361 |
| GPi/GPe | 0.0738 | ns | No | Two-tailed | t=2.163, df=6 | 4.677, 1, 6 | -0.1089 ± 0.05035 | 0.9532 ± 0.356 |
| MH | 0.079 | ns | No | Two-tailed | t=2.344, df=4 | 5.494, 1, 4 | -0.1204 ± 0.05138 | 1.371 ± 0.261 |
| VI | 0.0989 | ns | No | Two-tailed | t=1.867, df=8 | 3.484, 1, 8 | -0.09793 ± 0.05247 | 0.6598 ± 0.24 |
| NAc | 0.1108 | ns | No | Two-tailed | t=1.749, df=10 | 3.060, 1, 10 | -0.03805 ± 0.02175 | 0.486 ± 0.214 |
| MS | 0.3346 | ns | No | Two-tailed | t=1.067, df=5 | 1.139, 1, 5 | -0.04699 ± 0.04403 | 0.2562 ± 0.307 |
| OT | 0.7205 | ns | No | Two-tailed | t=0.3751, df=6 | 0.1407, 1, 6 | -0.02020 ± 0.05387 | 0.022 ± 0.369 |
| BW: Non-ChAT(M) v ChAT(M) | 0.0099 | ** | Yes | Two-tailed | t=5.862, df=3 | 34.36, 1, 3 | -0.1287 ± 0.02196 | |
| BW: Non-ChAT(F) v ChAT(F) | 0.0106 | * | Yes | Two-tailed | t=4.533, df=4 | 20.54, 1, 4 | -0.1208 ± 0.02664 | |
| BW: ChAT(F) v ChAT(M) | 0.437 | ns | No | Two-tailed | t=0.8945, df=3 | 0.8001, 1, 3 | -0.02941 ± 0.03288 | |
| BW: Non-ChAT(F) v Non-ChAT(M) | 0.3357 | ns | No | Two-tailed | t=1.093, df=4 | 1.195, 1, 4 | -0.02104 ± 0.01925 | |

**Figure 4 — Supplement 1**. CNS-wide Atlas of SPOTlight Activity in ChAT-positive neurons. (**A**) Statistical summary table of R:G ratio values of regional ChAT vs non-ChAT neurons related to **Figure 4 A-B**. The underlying data between R:G ratios and $log_2$ fold-change are the same with the same statistical summaries with the exception of mean differences. (**B**) Relative RFP fluorescence used to determine R:G ratios used in **Figure 4 A-B** with statistical summary table of differences in RFP abundance. (**C**) Relative GFP fluorescence used to determine R:G ratios used in **Figure 4 A-B** with statistical summary table of differences in GFP abundance.

**Figure 4 — Supplement 1** (continued)

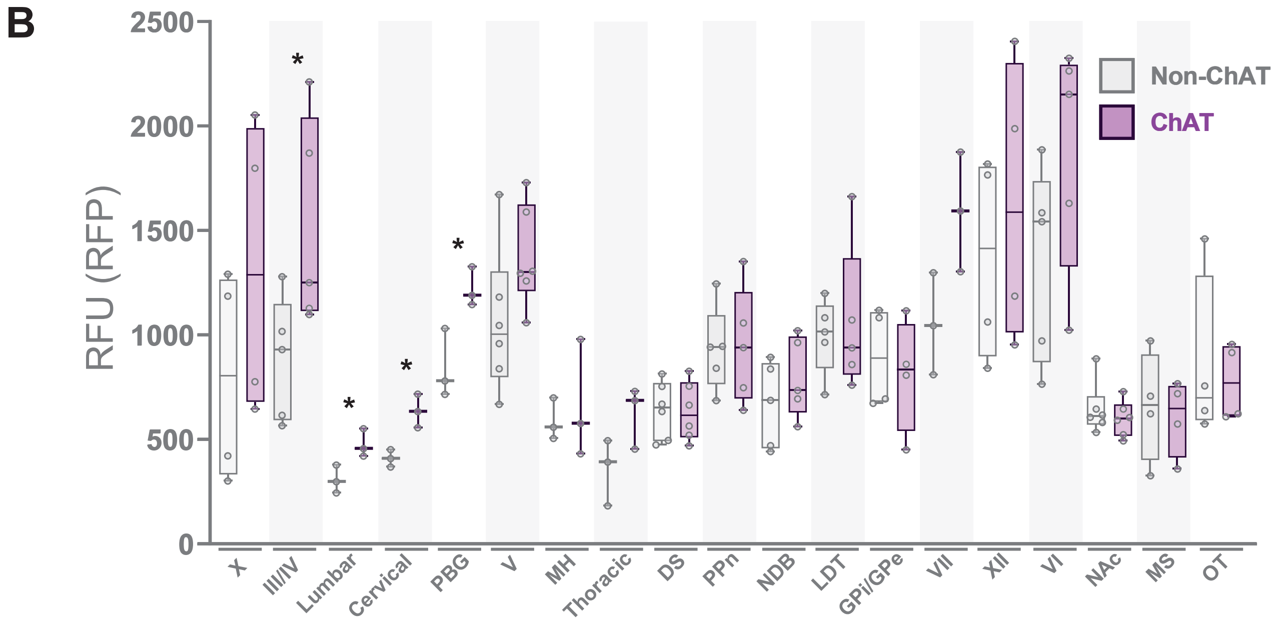


**Figure 4 Statistical Summary (Nested t Test: RFP Relative Abundance)**

| Non-ChAT vs ChAT | P value | P value summary | P < 0.05? | One- or two-tailed P value? | t, df | F, DFn, Dfd | Mean difference ± SEM |
|---|---|---|---|---|---|---|---|
| Cervical | 0.0129 | * | Yes | Two-tailed | t=4.276, df=4 | 18.28, 1, 4 | -226.3 ± 52.92 |
| PBG | 0.0226 | * | Yes | Two-tailed | t=3.609, df=4 | 13.03, 1, 4 | -385.4 ± 106.8 |
| Lumbar - no outlier | 0.0383 | * | Yes | Two-tailed | t=3.044, df=4 | 9.265, 1, 4 | -168.7 ± 55.41 |
| III/IV | 0.0419 | * | Yes | Two-tailed | t=2.420, df=8 | 5.856, 1, 8 | -630.4 ± 260.5 |
| VII | 0.0666 | ns | No | Two-tailed | t=2.502, df=4 | 6.262, 1, 4 | -543.5 ± 217.2 |
| Thoracic | 0.0877 | ns | No | Two-tailed | t=2.249, df=4 | 5.058, 1, 4 | -286.2 ± 127.3 |
| V | 0.1037 | ns | No | Two-tailed | t=1.790, df=10 | 3.205, 1, 10 | -312.7 ± 174.7 |
| VI | 0.14 | ns | No | Two-tailed | t=1.638, df=8 | 2.684, 1, 8 | -527.6 ± 322.1 |
| X | 0.2824 | ns | No | Two-tailed | t=1.181, df=6 | 1.394, 1, 6 | -515.8 ± 436.8 |
| NDB | 0.3458 | ns | No | Two-tailed | t=1.002, df=8 | 1.004, 1, 8 | -128.8 ± 128.5 |
| NAc | 0.4495 | ns | No | Two-tailed | t=0.7870, df=10 | 0.6194, 1, 10 | 46.98 ± 59.70 |
| XII | 0.5587 | ns | No | Two-tailed | t=0.6189, df=6 | 0.3830, 1, 6 | -260.2 ± 420.4 |
| GPi/GPe | 0.6774 | ns | No | Two-tailed | t=0.4370, df=6 | 0.1910, 1, 6 | 79.69 ± 182.3 |
| MH | 0.6902 | ns | No | Two-tailed | t=0.4287, df=4 | 0.1838, 1, 4 | -74.57 ± 173.9 |
| OT | 0.6976 | ns | No | Two-tailed | t=0.4078, df=6 | 0.1663, 1, 6 | 97.63 ± 239.4 |
| LDT | 0.7275 | ns | No | Two-tailed | t=0.3609, df=8 | 0.1303, 1, 8 | -64.52 ± 178.8 |
| MS | 0.7597 | ns | No | Two-tailed | t=0.3202, df=6 | 0.1025, 1, 6 | 52.00 ± 162.4 |
| PPn | 0.9082 | ns | No | Two-tailed | t=0.1190, df=8 | 0.01416, 1, 8 | -18.35 ± 154.2 |
| DS | 0.9249 | ns | No | Two-tailed | t=0.09666, df=10 | 0.0093, 1, 10 | 7.933 ± 82.07 |

### **Figure 4 — Supplement 1** (continued)

C

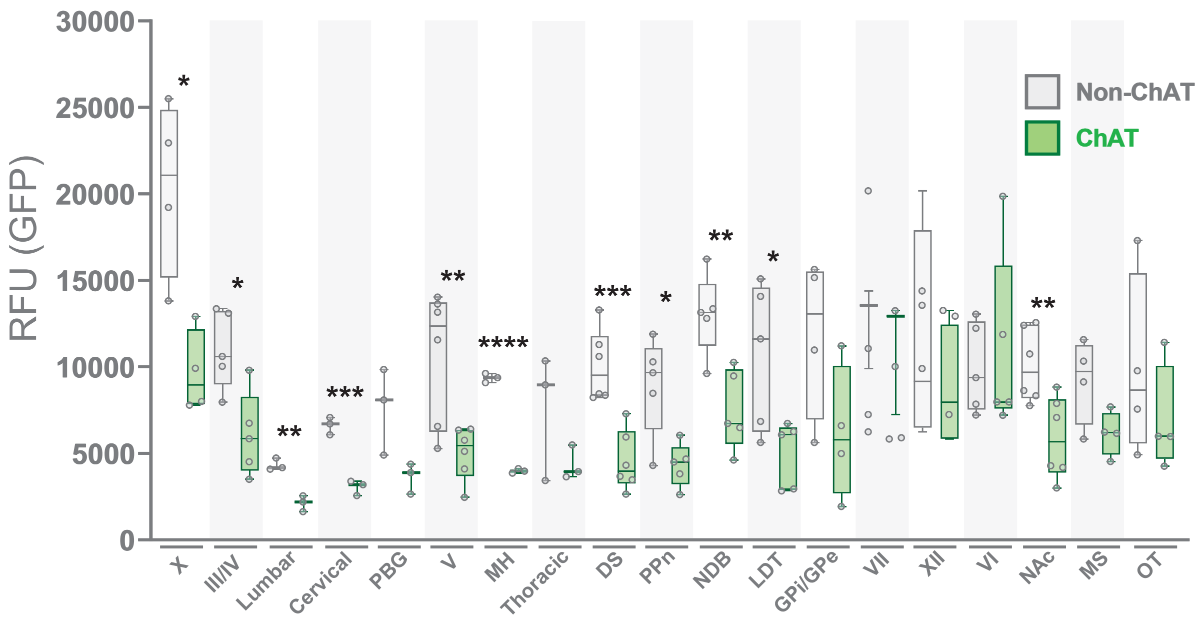


| Figure 4 Statistical Summary (Nested t Test: GFP Relative Abundance) | | | | | | | |
|---|---|---|---|---|---|---|---|
| **Non-ChAT vs ChAT** | **P value** | **P value summary** | **P < 0.05?** | **One- or two-tailed P value?** | **t, df** | **F, DFn , Dfd** | **Mean difference ± SEM** |
| MH | <0.0001 | **** | Yes | Two-tailed | t=28.75, df=4 | 826.3, 1, 4 | 5267 ± 183.2 |
| DS | 0.0006 | *** | Yes | Two-tailed | t=4.980, df=10 | 24.80, 1, 10 | 5480 ± 1100 |
| Cervical | 0.0007 | *** | Yes | Two-tailed | t=9.313, df=4 | 86.74, 1, 4 | 3550 ± 381.2 |
| Lumbar - no outlier | 0.0025 | ** | Yes | Two-tailed | t=6.725, df=4 | 45.23, 1, 4 | 2203 ± 327.6 |
| NDB | 0.0055 | ** | Yes | Two-tailed | t=3.760, df=8 | 14.14, 1, 8 | 5648 ± 1502 |
| V | 0.0064 | ** | Yes | Two-tailed | t=3.433, df=10 | 11.78, 1, 10 | 5722 ± 1667 |
| NAc | 0.0089 | ** | Yes | Two-tailed | t=3.240, df=10 | 10.50, 1, 10 | 4190 ± 1293 |
| III/IV | 0.0105 | * | Yes | Two-tailed | t=3.322, df=8 | 11.04, 1, 8 | 4915 ± 1479 |
| X | 0.0111 | * | Yes | Two-tailed | t=3.617, df=6 | 13.08, 1, 6 | 10494 ± 2901 |
| PPn | 0.0117 | * | Yes | Two-tailed | t=3.252, df=8 | 10.58, 1, 8 | 4583 ± 1409 |
| LDT | 0.0259 | * | Yes | Two-tailed | t=2.728, df=8 | 7.444, 1, 8 | 5681 ± 2082 |
| PBG | 0.0607 | ns | No | Two-tailed | t=2.589, df=4 | 6.704, 1, 4 | 3983 ± 1538 |
| MS | 0.0708 | ns | No | Two-tailed | t=2.193, df=6 | 4.807, 1, 6 | 3074 ± 1402 |
| GPi/GPe | 0.1089 | ns | No | Two-tailed | t=1.882, df=6 | 3.542, 1, 6 | 5678 ± 3017 |
| Thoracic | 0.214 | ns | No | Two-tailed | t=1.476, df=4 | 2.179, 1, 4 | 3217 ± 2179 |
| OT | 0.3753 | ns | No | Two-tailed | t=0.9576, df=6 | 0.9169, 1, 6 | 2986 ± 3119 |
| XII | 0.5246 | ns | No | Two-tailed | t=0.6754, df=6 | 0.4562, 1, 6 | 2477 ± 3667 |
| VII | 0.5885 | ns | No | Two-tailed | t=0.5873, df=4 | 0.3450, 1, 4 | 1428 ± 2432 |
| VI | 0.69 | ns | No | Two-tailed | t=0.4136, df=8 | 0.1711, 1, 8 | -1098 ± 2655 |

**Figure 4 — Supplement 2.** Continuation of representative images from CNS-wide Atlas of SPOTlight Activity in ChAT-positive neurons. (**A**) Medial septal nucleus (MS), scale bar 200 μm. (**B**) Olfactory tubercle (OT), scale bar 100 μm. (**C**) Diagonal band nucleus (NDB), scale bar 100 μm. (**D**) Globus pallidus external and internal segment (GPe/GPi), scale bar 50 μm. (**E**) Medial habenula (MH), scale bar 50 μm. (**F**) Parabigeminal nucleus (PBG), scale bar 50 μm. (**G**) Abducens nucleus (VI), scale bar 50 μm. (**H**) Motor nucleus of trigeminal (V) and Facial motor nucleus (VII), scale bar 200 μm. (**E-H**) continued on next page.

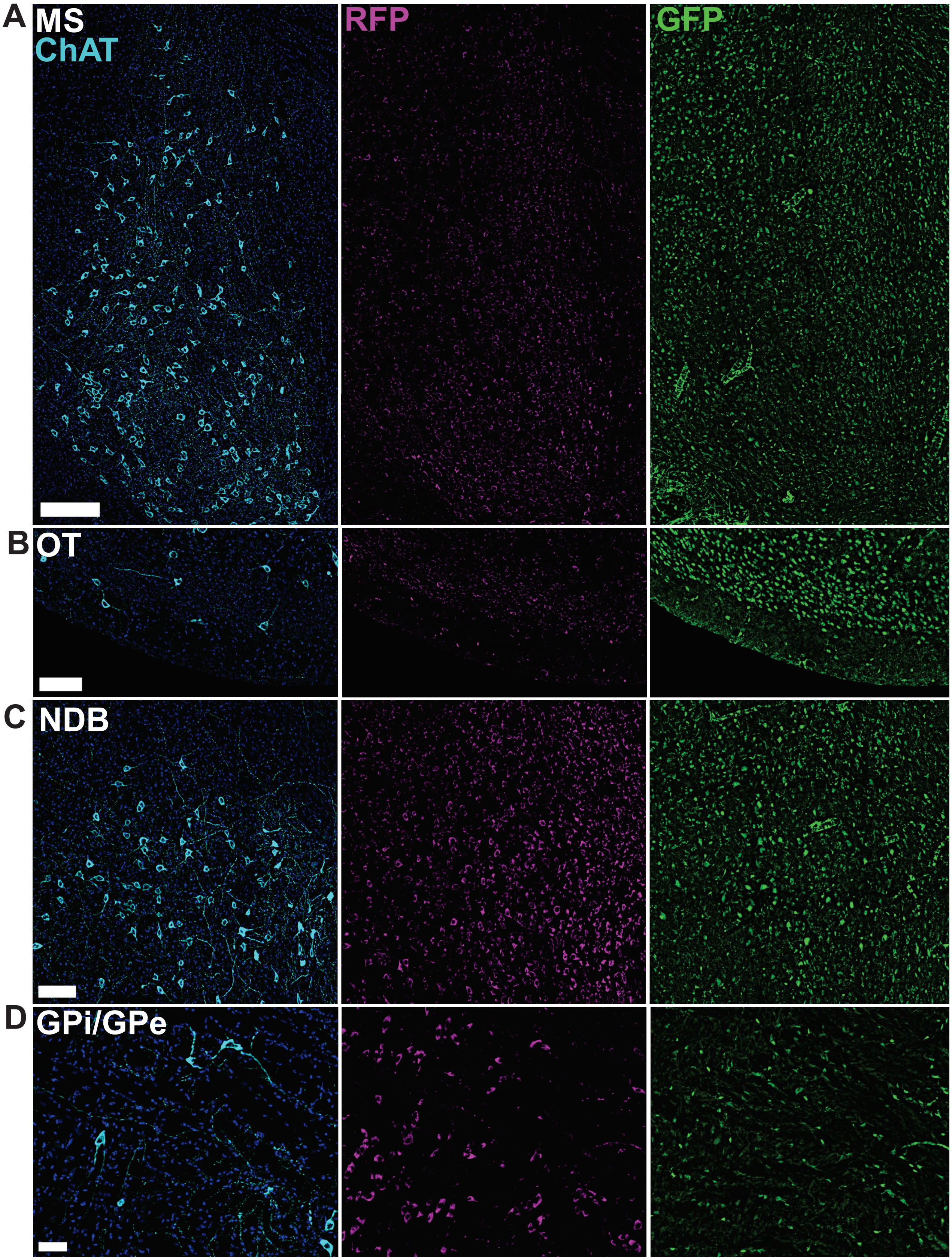

Figure 4 — Supplement 2 (continued)

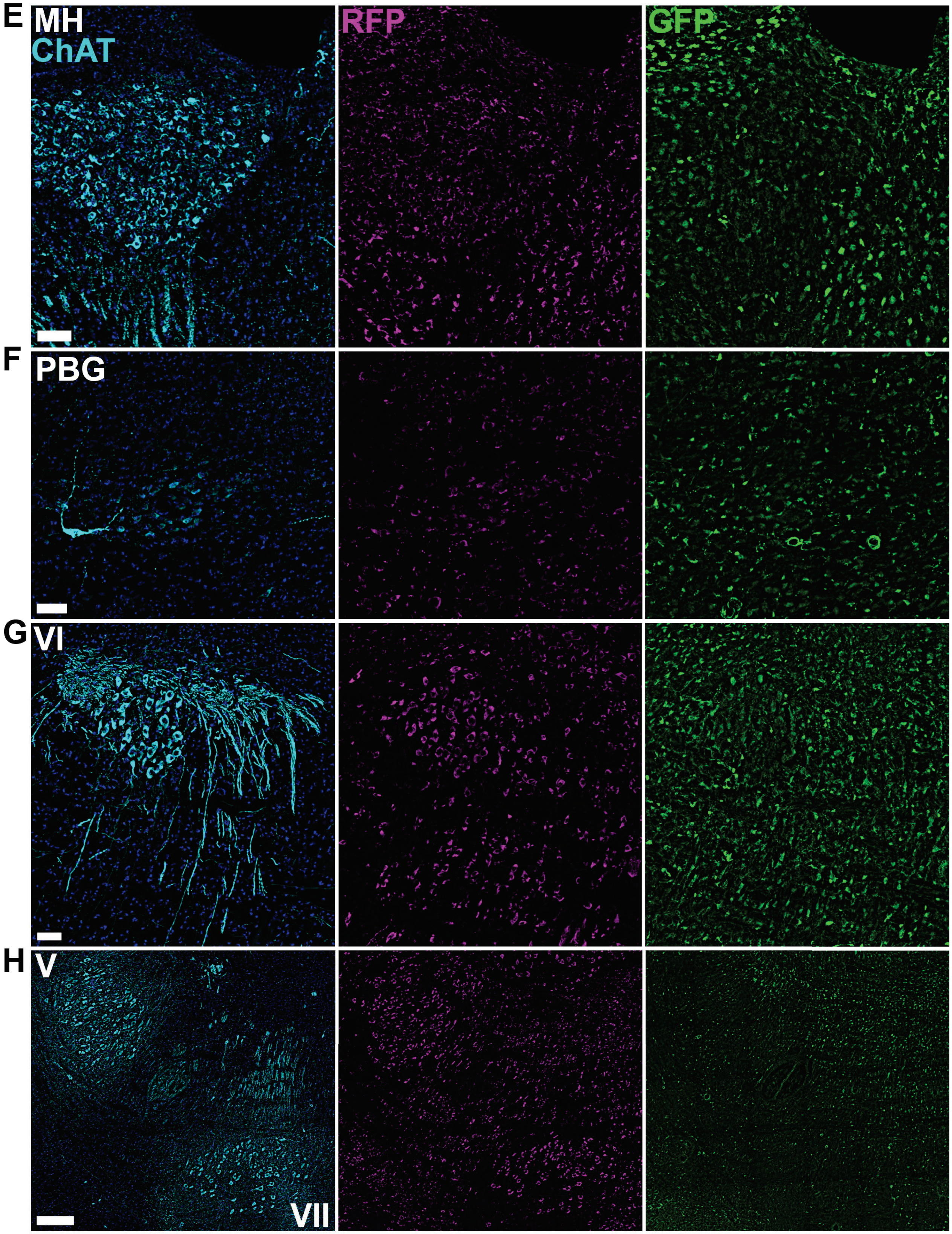
